# Development of a Wearable Sensor-PAM for Continuous Monitoring of Photosynthetic Dynamics

**DOI:** 10.64898/2026.09.12.751108

**Authors:** Ko-ichiro Miyamoto, Kentaro Ifuku, Tatsuo Yoshinobu, Kaori Kohzuma

**Affiliations:** Department of Electronic Engineering, Tohoku University, Sendai, Miyagi 980-8579, Japan; Division of Applied Life Sciences, Graduate School of Agriculture, Kyoto University, Kyoto 606-8502, Japan; Graduate School of Biomedical Engineering, Tohoku University, Sendai, Miyagi 980-8579, Japan

**Keywords:** Wearable sensor, Plant monitoring, Remote sensing, Environmental response, pulse-amplitude modulation (PAM) fluorometers

## Abstract

Continuous monitoring of photosynthetic performance is essential for understanding plant responses to fluctuating environments and has important applications in plant physiology, field phenotyping, and digital agriculture. However, conventional pulse-amplitude modulation (PAM) fluorometers are primarily designed for point measurements and are not suitable for long-term monitoring while attached to intact leaves. Here, we developed Sensor-PAM, a wearable chlorophyll fluorescence measurement system capable of continuously monitoring photosynthetic dynamics from the abaxial side of a leaf. The system combines a commercially available color sensor with blue LEDs in a compact, low-cost optical design to perform PAM measurements. Chlorophyll fluorescence measured from the abaxial leaf surface showed a strong correlation with conventional adaxial measurements and accurately reflected changes in photosynthetic performance induced by chilling and high-light stress. Measurements obtained using Sensor-PAM also showed good agreement with those from a commercial PAM fluorometer across diverse plant species. Furthermore, the wearable system enabled continuous monitoring of the effective quantum yield of photosystem II [Y(II)] from the same position on the same strawberry leaf for three days under both greenhouse and outdoor conditions, successfully capturing photosynthetic responses to changing irradiance and temperature in real time. These findings establish Sensor-PAM as a wearable platform for continuous chlorophyll fluorescence monitoring, extending conventional PAM fluorometry from point-based measurements to long-term monitoring of photosynthetic dynamics under natural environmental conditions.

## 1. Introduction

Plants continuously adjust their photosynthetic activity in response to fluctuating environmental conditions, including changes in light intensity, temperature, water availability, and nutrient status (Li et al., 2023; Vialet-Chabrand et al., 2017). Photosynthetic impairment caused by environmental stress often occurs before visible symptoms, such as leaf discoloration or wilting, become apparent (Baker, 2008; Maxwell and Johnson, 2000). Therefore, non-destructive and temporal monitoring of photosynthetic performance has become increasingly important not only in fundamental plant physiology but also in agriculture, plant breeding, ecology, and digital farming (Araus and Cairns, 2014; Chawade et al., 2019; Kohzuma et al., 2021; Kuhlgert et al., 2016). Among the available approaches, chlorophyll fluorescence is widely used as a sensitive indicator of photosystem II (PSII) activity and plant responses to environmental stress (Murchie and Lawson, 2013; Swoczyna et al., 2022).

Pulse-amplitude modulation (PAM) fluorometry is a standard method for quantitative chlorophyll fluorescence analysis. By combining weak measuring light with saturating light pulses, PAM fluorometry enables the non-destructive determination of minimum fluorescence (Fo), maximum fluorescence (Fm), the maximum quantum yield of PSII (Fv/Fm), and the effective quantum yield of PSII (Y(II)) (Genty et al., 1989; Kalaji et al., 2017; Schreiber et al., 1986). These parameters sensitively detect changes in PSII function caused by photoinhibition, chilling, drought, and other environmental stresses, making PAM fluorometry one of the most widely used approaches for evaluating photosynthetic performance (Baker, 2008; Stirbet et al., 2018). Despite its widespread use, commercial PAM fluorometers are generally expensive, rely on complex optical systems, and often require manual positioning of a probe on the leaf, making them unsuitable for long-term continuous monitoring (Haidekker et al., 2022).

The Monitoring-PAM fluorometer (Walz, Germany) was developed to enable long-term continuous chlorophyll fluorescence measurements under field conditions (Porcar - Castell, 2011; Zhang et al., 2025). Although this represented a major advance, the system remains expensive, and the placement of the sensor on the adaxial leaf surface may shade the leaf and alter its natural light environment during measurement. More recently, several low-cost and open-source chlorophyll fluorometers have been developed, including Open-JIP, an Arduino-based low-cost OJIP fluorometer (Bates et al., 2019), a PAM fluorometer that can be assembled for less than 300 US dollars (Haidekker et al., 2022), an autonomous wireless PAM fluorometer costing approximately 170 euros (Baghbani et al., 2026) and the Active Pulsed System (APS) designed for long-term monitoring (Astashev et al., 2026). Simplified chlorophyll fluorescence imaging systems that do not rely on PAM have also been reported for plant stress detection (Legendre et al., 2021). These developments have improved the affordability, automation, and field applicability of chlorophyll fluorescence measurements. Nevertheless, most existing systems are either handheld or fixed on a leaf and are not designed for continuous measurements while attached to intact leaves. In addition, placing a sensor on or above the adaxial leaf surface may interfere with incident light and thereby alter the local light environment at the measurement site. This problem can potentially be avoided by measuring chlorophyll fluorescence from the abaxial surface; however, fluorescence characteristics measured from the adaxial and abaxial surfaces are not necessarily identical because of differences in leaf anatomy and the absorption and scattering of light within the leaf (Cordon and Lagorio, 2007; Terashima and Saeki, 1983; Vogelmann and Evans, 2002). Therefore, before abaxial measurements can be applied to continuous monitoring under natural light conditions, it is necessary to determine whether chlorophyll fluorescence measured from the abaxial surface reliably reflects changes in photosynthetic performance.

In parallel with advances in chlorophyll fluorescence instrumentation, wearable plant sensors that can be directly attached to leaves have rapidly emerged as promising tools for continuous plant monitoring. Various devices have been developed to monitor leaf temperature, water status, growth, volatile organic compounds, and other plant traits, with potential applications in crop diagnosis and digital agriculture (Lee et al., 2024; Nassar et al., 2018; Yin and Dong, 2024). We previously developed a compact, low-cost wireless plant sensor that can be attached to the abaxial leaf surface and continuously acquire reflectance spectra without blocking incident light on the adaxial surface (Kohzuma and Miyamoto, 2024). The sensor transmitted data to cloud storage via Wi-Fi, enabling real-time remote monitoring of plant status. Using this system, we detected changes in chlorophyll content and reflectance associated with the xanthophyll cycle (Blackburn, 1998; Gamon et al., 1992; Gitelson et al., 2003). Although reflectance measurements provide useful information on plant physiological status, they do not directly quantify photosynthetic function and have limited ability to detect rapid changes in PSII activity under fluctuating light and temperature conditions (Murchie and Lawson, 2013).

To overcome this limitation, we integrated PAM fluorometry into our previously developed wearable sensor platform and developed a wearable pulse-amplitude modulation chlorophyll fluorometer, termed Sensor-PAM (Fig. 1). The use of a commercially available color sensor was central to the miniaturization of the system because the sensor integrates photodetectors, current-detection circuitry, an analog-to-digital converter, and a communication interface into a single chip. This architecture enabled the construction of a compact sensor head that can be attached to the abaxial leaf surface while preserving the natural light environment of the adaxial surface.

**Fig. 1.**
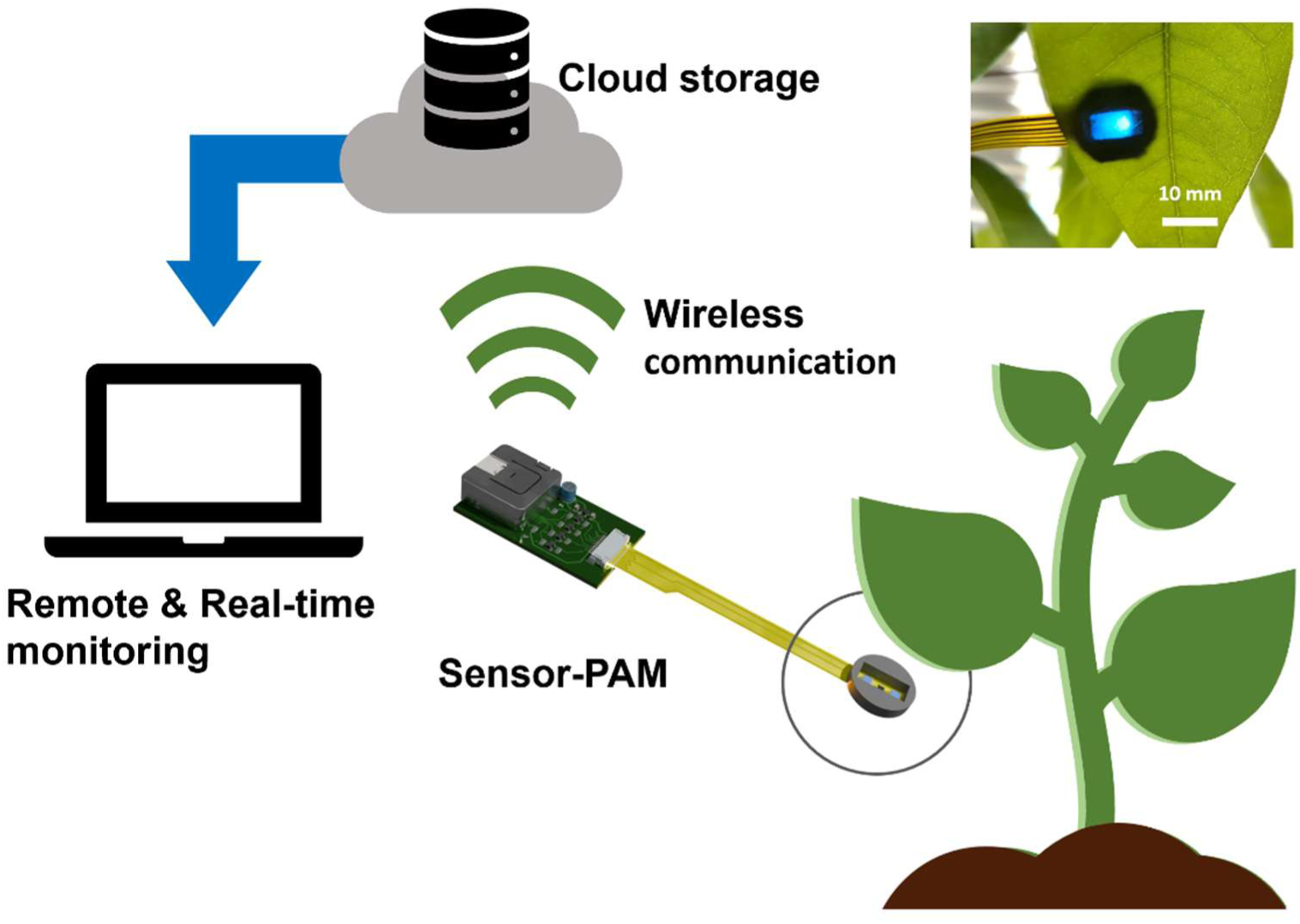
Concept of the wearable Sensor-PAM for continuous chlorophyll fluorescence monitoring. The sensor is attached to the abaxial leaf surface, and measurement data are wirelessly transmitted to a cloud platform for remote, real-time monitoring.

In this study, we first examined whether chlorophyll fluorescence measured from the abaxial leaf surface could evaluate photosynthetic status comparably to conventional measurements from the adaxial surface. We then validated the performance of the Sensor-PAM by comparison with a commercial PAM fluorometer across multiple plant species and under field conditions. Finally, we developed Wearable Sensor-PAM, a wearable version of the system and demonstrated continuous monitoring of photosynthetic dynamics under greenhouse and outdoor conditions.

## 2. Material and methods

### 2.1 Sensor-PAM

A prototype Sensor-PAM system used for the evaluation is depicted in Fig. 2A. It consisted of a color sensor, two LEDs, and a controller connected to a PC. The color sensor (AS7341, ams-OSRAM AG, Premstaetten, Austria) was implemented on a breakout board (SEN0365, DFRobot, Pudong New Area, China). Two blue LEDs (λ = 470 nm) were used to provide the measuring light (ML) and saturation flash (SF), respectively. The ML was used to monitor the fluorescence signal, whereas the SF was applied to induce maximum fluorescence. The intensities of the ML and SF were adjustable. These components, along with a homemade spacer designed to maintain a constant distance between the sensor head and the leaf surface to improve measurement reproducibility (Fig. 2B), were mounted and secured onto a custom-designed printed-circuit board (PCB, thickness = 1.5 mm).

**Fig. 2.**
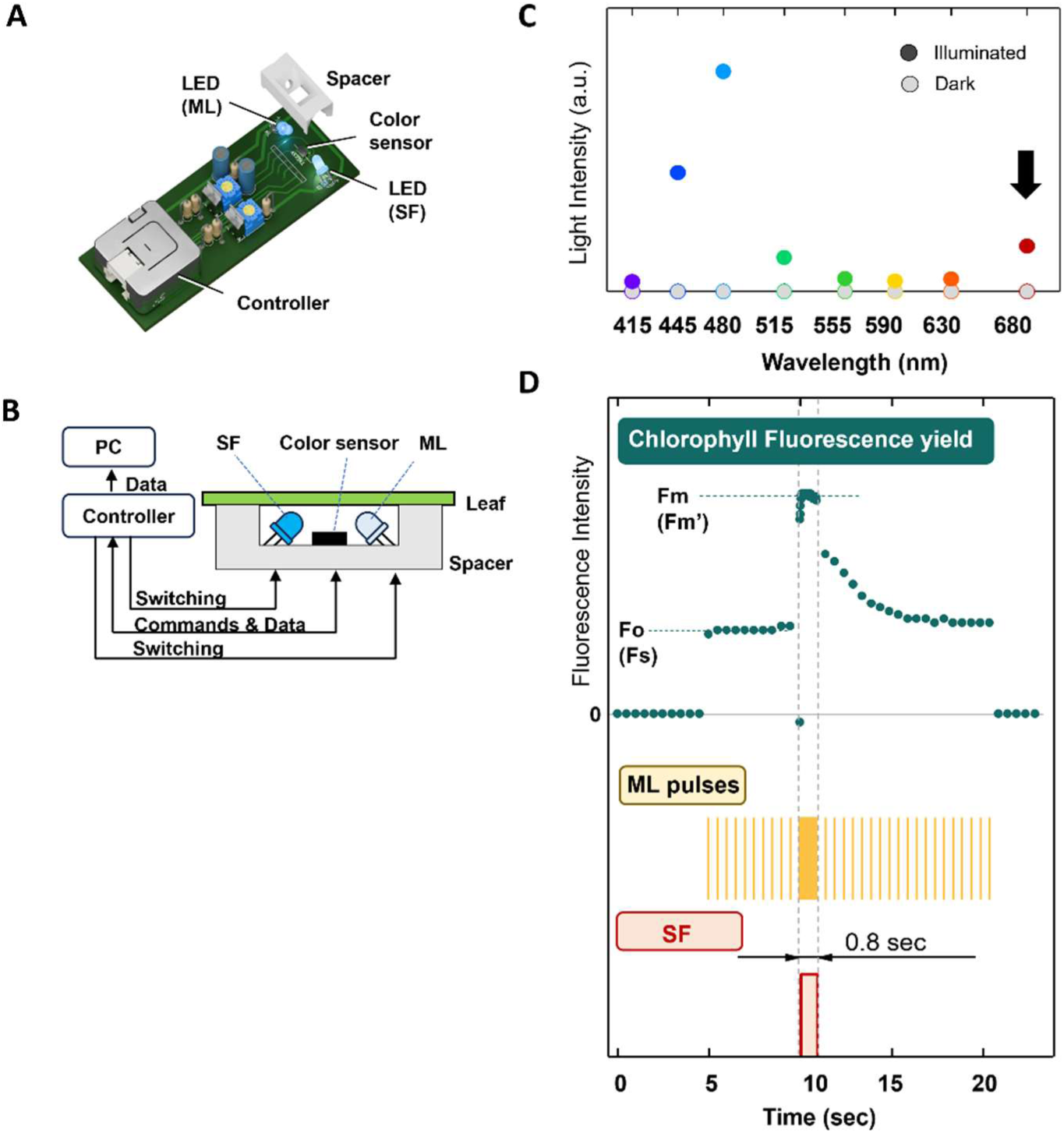
Prototype Sensor-PAM and its operating principle. (A) Image of the prototype Sensor-PAM. (B) Schematic illustration of the Sensor-PAM. A blue LED provides measuring light (ML) and a saturating flash (SF), while chlorophyll fluorescence emitted from the abaxial side of the leaf is detected by the integrated color sensor. The Sensor-PAM is controlled by a microcontroller and communicates with a personal computer for data acquisition and instrument control. (C) Detection of chlorophyll fluorescence using the eight-channel color sensor. Gray circles represent sensor signals measured in the dark, whereas color-filled circles represent signals measured during a blue saturating pulse. The signals at445–480nm correspond primarily to the reflected blue excitation light, whereas the signal at 680 nm represents chlorophyll fluorescence. (D) Measurement sequence of the Sensor-PAM. Weak ML pulses was applied to monitor chlorophyll fluorescence, and SF pulses were applied to induce the maximum fluorescence yield (Fm in dark-adapted measurements or Fm’ in light-adapted measurements). The fluorescence level immediately before the saturating pulse was recorded as Fo or Fs, respectively.

Prior to the development of the Sensor-PAM, the spectral response of the color sensor was examined to select the appropriate channel for chlorophyll fluorescence detection. Each channel of the color sensor covers a relatively broad wavelength range of several tens of nanometers because of the laminated filters integrated into the sensor. Fig. 2C shows a typical response of the color sensor to reflected excitation light and chlorophyll fluorescence from a leaf. Under dark conditions, minimal signal was detected in any wavelength channel. Following application of a blue saturation flash, the signal intensity increased at 445–480 nm, corresponding primarily to reflected excitation light, whereas a pronounced increase was observed in the 680-nm channel, corresponding to chlorophyll fluorescence. These results indicated that the 680-nm channel was suitable for chlorophyll fluorescence detection. To prevent the detection of stray light, the sensor was covered with a red optical film in the following experiments to ensure separation between excitation light and fluorescence.

### 2.2 Measurement software

A measurement program for the controller (M5 Atom Lite, M5Stack Technology Co., Ltd, Shenzhen, China) was developed using the Arduino IDE (Arduino LLC, Monza, Italy). The program operated both the LEDs and the color sensor. The data were transferred from the controller to the PC via serial communication.

Fig. 2D shows an example of the fluorescence signal obtained using the measurement program. Steady-state fluorescence was monitored using measuring light, and maximum fluorescence (Fm) was obtained by applying a saturation flash. A typical PAM fluorescence trace was successfully recorded, allowing stable determination of both minimum fluorescence (Fo) and maximum fluorescence (Fm). To obtain the accurate values for the fluorescence yield, especially the Fo obtained under dark conditions, it is essential to keep the ML as short as possible to avoid the actinic effect on photosynthesis. In this study, the measurement program set the illumination time of the ML to 75 µs, which was equal to the integration time of the color sensor. The interval between the ML pulses was set to 0.5 s, but was reduced to 150 µs only during the SF irradiation (0.8 s) to capture the detailed response of the fluorescence yield. The program was designed to synchronize these operations of the LEDs and data collection. To minimize the processing overhead during this synchronization, only the 680 nm channel was read out in the measurement program.

### 2.3 Wearable Sensor-PAM

Based on the evaluation of the prototype system, the wearable version of the Sensor-PAM was implemented. To minimize the size and weight of the sensor head, the sensor system was divided into an interface section for the controller and a sensor head section, which were fabricated on a rigid PCB and a flexible printed circuit (FPC), respectively. The driver circuits for the LEDs and the color sensor were implemented in the interface section, while the sensor head consisted of only two surface-mounted LEDs and the color sensor. The sensor head measures 8 × 175 mm in size, weighing 0.45 g, and remains as light as 1.8 g with a ring-shaped spacer. The sensor head was attached to the abaxial surface of the leaf using double-sided tape. When operating the wearable Sensor-PAM, the data were transferred via Wi-Fi communication to cloud storage.

### 2.4 Measurement of Y(II)

In this study, chlorophyll fluorescence measurements were performed primarily under light-adapted conditions. The Sensor-PAM was developed not only to evaluate the maximum quantum efficiency of PSII after dark adaptation but also to monitor changes in PSII photochemical efficiency under natural light environments and environmental stress conditions. Therefore, in most experiments, leaves were measured without prior dark adaptation, and a saturating pulse was applied to leaves in their ambient light-adapted state. The steady-state fluorescence immediately before the saturating pulse and the maximum fluorescence induced by the saturating pulse were defined as Fs and Fm′, respectively. The effective quantum yield of PSII, Y(II), was calculated as:

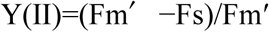

In some experiments (e.g., Fig. 3A-a), leaves were dark-adapted prior to the measurement, and the resulting values corresponded to the conventional maximum quantum efficiency of PSII (Fv/Fm). However, for consistency throughout this paper, these values are also referred to as Y(II).

**Fig. 3.**
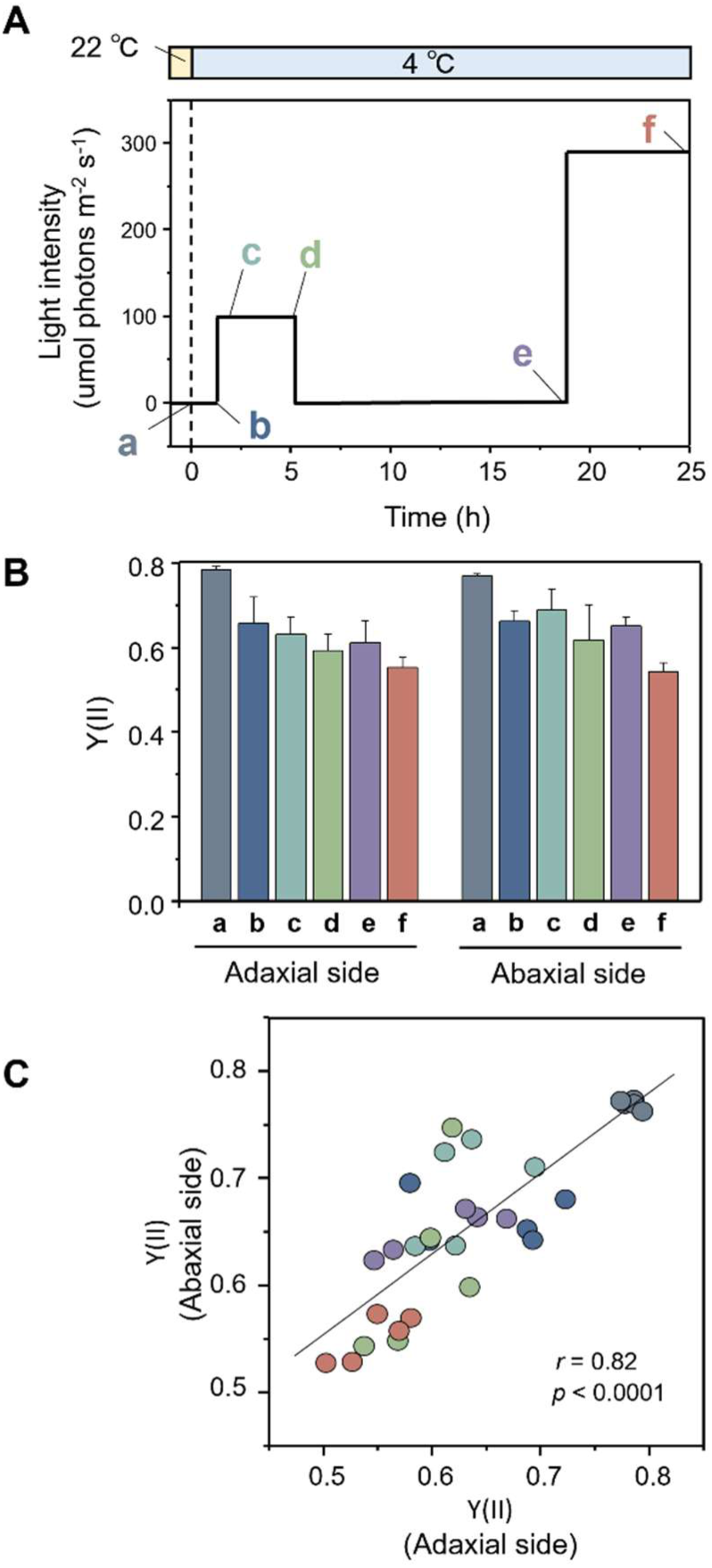
Validation of abaxial chlorophyll fluorescence measurements. (A) Schematic of the low-temperature/high-light treatment applied to *Arabidopsis thaliana*. After dark adaptation at 22°C (a, gray), plants were transferred to 4°C and kept in the dark for 1 h (b, blue). Leaves were then illuminated at 100 μmol m⁻² s⁻¹ for 1 h (c, teal) and 5 h (d, green), followed by 14 h of dark treatment at 4°C (e, purple). Finally, leaves were exposed to high light (200 μmol m⁻² s⁻¹) for 5 h at 4°C (f, reddish brown). (B) Effective quantum yield of PSII (Y(II)) measured from the adaxial and abaxial sides of the same leaves using a reference PAM fluorometer. Data are presented as mean ± SD (n = 5). (C) Correlation between Y(II) values measured from the adaxial and abaxial sides. The x-axis represents Y(II) measured from the adaxial side, and the y-axis represents Y(II) measured from the abaxial side.

### 2.5 Field experiment by the Sensor-PAM or the Wearable Sensor-PAM

The performance of the Sensor-PAM was evaluated under cold, sunny outdoor conditions between December 2025 and January 2026. Measurements were conducted on five days at two locations, Bunkyo-ku (Tokyo) and Sakyo-ku (Kyoto), Japan, between 09:00 and 15:00. A total of 101 leaves representing diverse plant species and light environments were examined, including herbaceous and woody plants, angiosperms and gymnosperms, monocots and dicots, and ferns (Fig. 5A).

To validate the performance of the Sensor-PAM under field conditions, chlorophyll fluorescence measurements were compared with those obtained using a commercially available portable PAM fluorometer (Junior-PAM; Walz, Germany). Measurements were performed under light-adapted conditions without prior dark adaptation, as described above. The fiber-optic probe of the Junior-PAM was secured in a leaf clip and positioned against the abaxial surface of the leaf, after which a single saturating pulse was applied. The Sensor-PAM head was then positioned at an adjacent location on the abaxial surface of the same leaf, and a single saturating pulse was applied in the same manner (Fig. 5B). The chlorophyll fluorescence parameter (Y(II)) obtained with the two instruments was subsequently compared.

The evaluation of the wearable Sensor-PAM was conducted in the winter season of February 2026 in Sendai, Japan (38.2555° N, 140.8412° E). The sensor head was attached to the abaxial surface of a potted strawberry leaf. Alongside the sample, both an illuminometer (LA-105, Nippon Medical & Chemical Instruments Co., Ltd, Japan) and a temperature logger (RC-5, Elitech Technology, Inc., USA) were installed to record the temporal changes. For the first three days, the measurements were conducted indoors (under greenhouse conditions). For the next three days, the sample was moved outdoors to monitor the response to low-temperature conditions.

## 3 Results

### 3.1 Validation of abaxial chlorophyll fluorescence measurements

The Sensor-PAM was designed to measure chlorophyll fluorescence from the abaxial leaf surface. Because the light environment and internal light propagation differ between the adaxial and abaxial surfaces, we first examined whether chlorophyll fluorescence measured from the abaxial side could reliably represent photosynthetic status. *Arabidopsis thaliana* plants were subjected to chilling and high-light treatments, and Y(II) was measured from both the adaxial and abaxial surfaces of the same leaves by using commercial PAM (Fig. 3A). On the adaxial surface, Y(II) progressively decreased during chilling and subsequent high-light treatment (Fig. 3B). A similar decrease was observed on the abaxial surface, although the reduction was slightly smaller. Comparison of Y(II) values obtained from both leaf surfaces revealed a significant positive correlation (Fig. 3C). A similarly strong correlation was also observed in strawberry leaves used in the experiments described in a later section (Fig. S1). These results indicate that, although chlorophyll fluorescence measurements from the abaxial and adaxial surfaces do not necessarily yield identical absolute values, abaxial measurements reliably capture changes in photosynthetic performance induced by chilling and high-light stress.

### 3.2 Validation of Sensor-PAM measurements

Having confirmed the validity of abaxial chlorophyll fluorescence measurements, we next evaluated the performance of the Sensor-PAM by comparing its measurements with those obtained using a commercial PAM fluorometer (Reference PAM fluorometer). *Arabidopsis thaliana* plants were exposed to chilling and high-light treatments, and Y(II) was measured from the abaxial leaf surface (Fig. 4A). Both the Reference PAM fluorometer and the Sensor-PAM detected similar decreases in Y(II) during stress treatment. The measurements obtained using the two systems were highly correlated (Fig. 4B), indicating that the Sensor-PAM accurately detected the decline in photosynthetic activity associated with photoinhibition. The same comparison was performed using strawberry leaves (Fig. 4C). Similar changes in Y(II) were observed with both instruments, and the measurements again showed a strong positive correlation (Fig. 4D). Although Y(II) values measured by the two instruments were strongly correlated, the regression slopes were less than 1 in both Arabidopsis and strawberry (Fig. 4B, D). These results demonstrate that the Sensor-PAM provides stable measurements across different plant species.

**Fig. 4.**
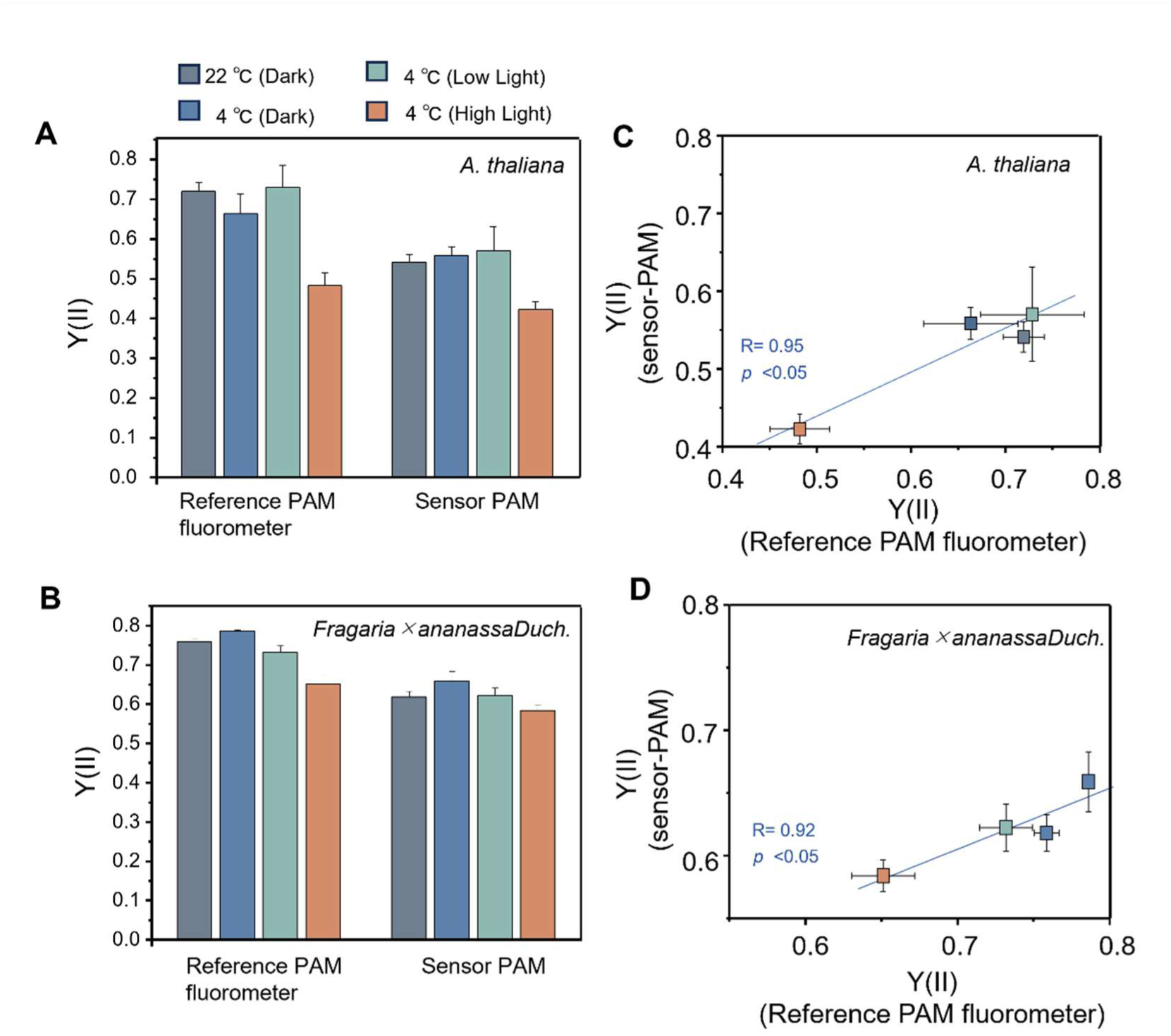
Validation of the Sensor-PAM for monitoring photoinhibition under low-temperature/high-light conditions. (A, C) Comparison of the effective quantum yield of PSII (Y(II)) measured using a reference PAM fluorometer (Junior-PAM, Walz) and the Sensor-PAM in *Arabidopsis thaliana* (A) and strawberry (*Fragaria × ananassa*) (C). After dark adaptation at 22°C (gray), leaves were transferred to 4°C and kept in the dark for 1 h (blue), illuminated at 100 μmol m⁻² s⁻¹ for 1 h (teal), and finally exposed to high light (500 μmol m⁻² s⁻¹) for 1 h at 4°C (reddish brown). Y(II) was measured from the abaxial side of the same leaves using both the reference PAM fluorometer and the Sensor-PAM. Data are presented as mean ± SD (n = 4). (B, D) Correlation between Y(II) values measured using the reference PAM fluorometer and the Sensor-PAM in *Arabidopsis thaliana* (B) and strawberry (D). Each point represents the mean Y(II) value obtained under each treatment condition. The x-axis represents Y(II) measured using the reference PAM fluorometer, and the y-axis represents Y(II) measured using the Sensor-PAM.

To further evaluate its applicability under natural conditions, field measurements were conducted over five days during winter using 101 leaves collected from diverse plant species (Fig. 5A, B). For each leaf, Y(II) was sequentially measured at the same position using both the Reference PAM fluorometer and the Sensor-PAM. Measurements obtained with the two systems showed a strong positive correlation (Fig. 5C). Bland– Altman analysis revealed that the Sensor-PAM systematically underestimated Y(II) by an average of 0.120 compared with the Reference PAM fluorometer (Fig. S2). The mean bias was −0.120, and the 95% limits of agreement ranged from −0.257 to 0.017. Despite this systematic offset, the high correlation and good agreement indicate that the Sensor-PAM provides reliable measurements across diverse plant species under natural environmental conditions.

**Fig. 5.**
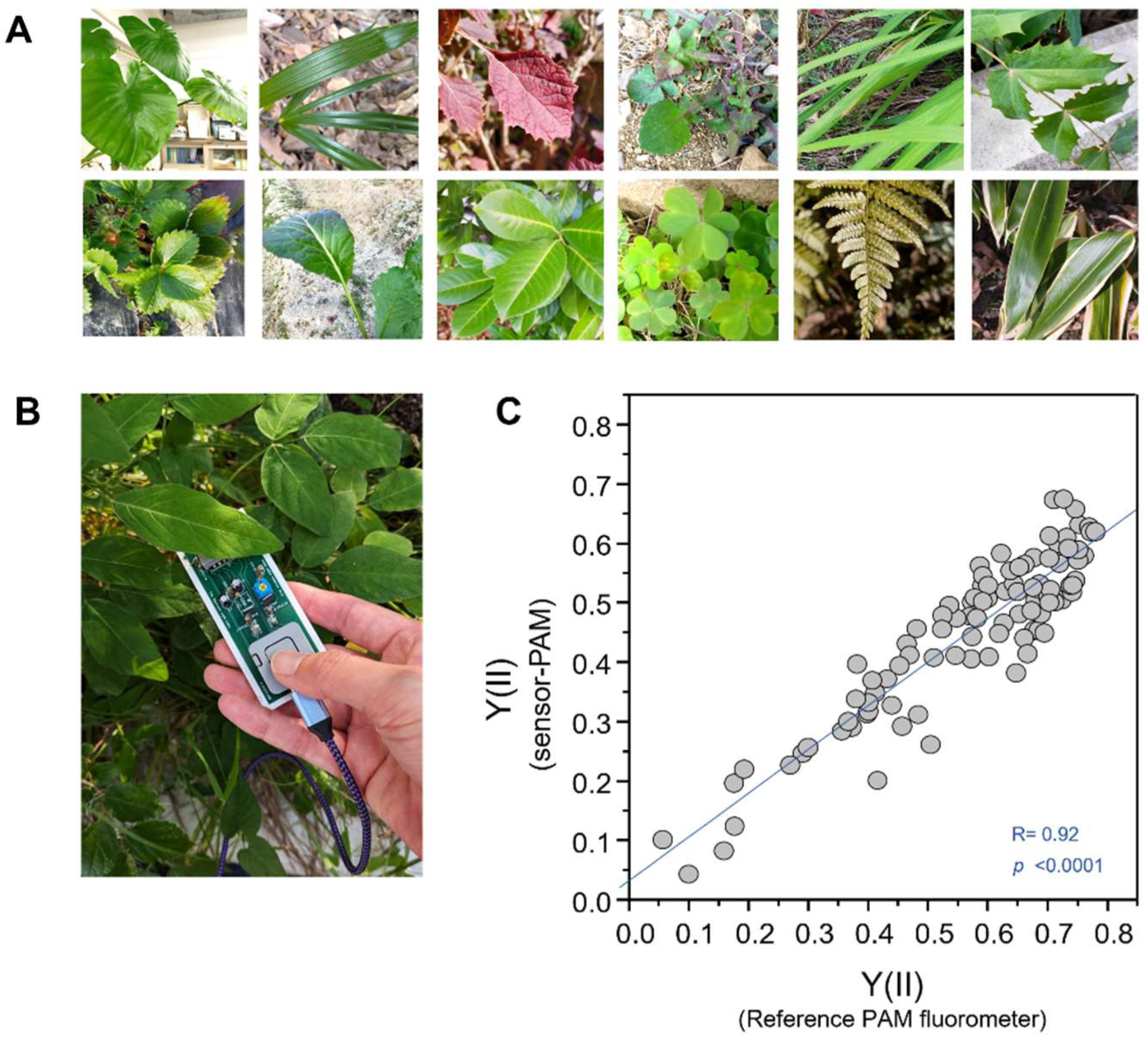
Field validation of the Sensor-PAM across diverse plant species. (A) Representative leaves used for field validation. Chlorophyll fluorescence measurements were performed on a total of 101 leaves collected from diverse plant species during five independent field surveys conducted under natural winter conditions. (B) Photograph of the field measurement. Chlorophyll fluorescence was measured non-destructively from the abaxial side of the same leaf at the same position using both the Sensor-PAM and a reference PAM fluorometer (Junior-PAM, Walz). (C) Correlation between Y(II) values measured using the reference PAM fluorometer and the Sensor-PAM. Measurements were performed on the abaxial side of 101 leaves. The x- and y-axes represent Y(II) measured using the reference PAM fluorometer and the Sensor-PAM, respectively.

### 3.3 Wearable Sensor-PAM enables continuous monitoring under natural conditions

Following validation of the prototype Sensor-PAM, we developed a wearable version capable of long-term measurements while attached directly to plant leaves (Fig. 6A-C). The wearable Sensor-PAM incorporates a compact and lightweight sensor head that can be stably attached to the abaxial leaf surface for continuous monitoring. The wearable Sensor-PAM was attached to strawberry leaves, and Y(II) was continuously monitored at the same position on the same leaf for three days under both greenhouse and outdoor conditions (Fig. 6D-E).

**Fig. 6.**
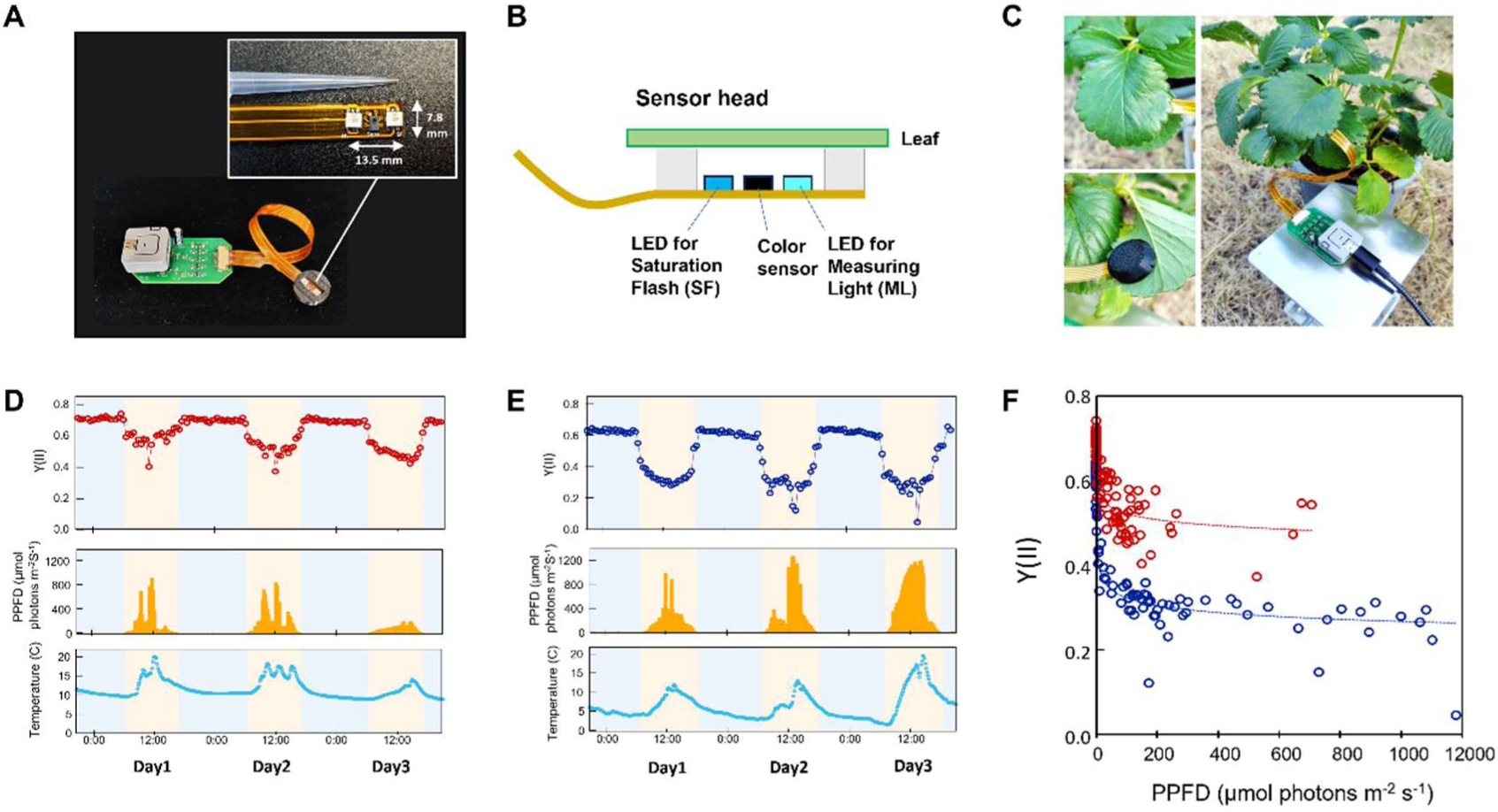
Continuous monitoring of photosynthetic dynamics using the wearable Sensor-PAM. (A) Photograph of the wearable Sensor-PAM. (B) Schematic cross-sectional view of the sensor head. Measuring light (ML) and saturation flash (SF) emitted from blue LEDs induce chlorophyll fluorescence, which is detected by a color sensor positioned between the LEDs. (C) Photographs showing the wearable Sensor-PAM attached to the abaxial side of a strawberry (*Fragaria × ananassa*) leaf and the experimental setup for continuous monitoring. The sensor was non-destructively attached to the abaxial leaf surface using a double-sided adhesive spacer, enabling repeated measurements at the same position. (D, E) Continuous monitoring of photosynthetic responses in strawberry leaves over three consecutive days under greenhouse (D) and outdoor (E) conditions. Red and blue traces represent the effective quantum yield of PSII (Y(II)) measured with the Sensor-PAM under greenhouse and outdoor conditions, respectively. Air temperature is shown as the light-blue line, and photosynthetic photon flux density (PPFD) is shown as yellow bars. Shaded areas indicate the light period. (F) Relationship between PPFD and Y(II) obtained from continuous measurements under greenhouse (red) and outdoor (blue) conditions. Solid lines indicate exponential regression curves.

Under greenhouse conditions, Y(II) exhibited regular diurnal fluctuations corresponding to the day–night cycle (Fig. 6D). In contrast, outdoor measurements showed not only diurnal variation but also rapid fluctuations in response to changes in irradiance and air temperature (Fig. 6E). These fluctuations closely corresponded to simultaneously recorded photosynthetic photon flux density (PPFD) and air temperature. Comparison of PPFD and Y(II) further showed that Y(II) decreased with increasing PPFD under both greenhouse and outdoor conditions (Fig. 6F). However, the magnitude of Y(II) showed substantially greater variation, reflecting the broader range of environmental fluctuations under natural conditions. Together, these results demonstrate that the wearable Sensor-PAM enables long-term continuous measurements at the same position on the same leaf, allowing real-time monitoring of photosynthetic dynamics under natural environmental conditions.

## 4. Discussion

### 4.1 Development of the wearable Sensor-PAM platform

Recent advances in low-cost chlorophyll fluorometers have substantially improved the affordability, automation, and field applicability of chlorophyll fluorescence measurements (Astashev et al., 2026; Baghbani et al., 2025; Bates et al., 2019; Haidekker et al., 2022). Nevertheless, most of these systems are either handheld or fixed instruments and are not intended for continuous measurements while attached to intact leaves.

The major innovation of the present study is the development of a wearable PAM fluorometer that can remain attached to a leaf throughout the measurement period. By fixing the sensor to the abaxial leaf surface, the Sensor-PAM measures chlorophyll fluorescence at the same position, thereby minimizing variation caused by probe positioning or spatial heterogeneity within the leaf (Perez-Bueno et al., 2019). This enables the photosynthetic dynamics of an individual leaf to be tracked directly over time. Continuous monitoring of Y(II) for three days under both greenhouse and outdoor conditions successfully captured regular diurnal fluctuations as well as rapid responses to changes in irradiance and temperature (Fig. 6). These findings demonstrate that the Sensor-PAM extends low-cost PAM fluorometry from portable measurements to continuous wearable monitoring of photosynthetic dynamics.

As a secondary benefit, utilizing a color sensor allowed for a simplified architecture, keeping the total component count to just 32. Consequently, the total cost of the wearable Sensor-PAM was reduced to approximately 30 USD (Table S1). This represents a cost advantage over previously reported systems (Baghbani et al., 2025; Haidekker et al., 2022), facilitating the deployment of multiple sensors for simultaneous and distributed monitoring.

### 4.2 Significance of abaxial chlorophyll fluorescence measurements

A key feature of the Sensor-PAM is that chlorophyll fluorescence is measured from the abaxial rather than the adaxial leaf surface. Previous studies have shown that chlorophyll fluorescence characteristics measured from the adaxial and abaxial surfaces are not necessarily identical. Within the leaf, the palisade and spongy mesophyll differ in their optical properties, and the absorption and scattering of excitation light, as well as the scattering and reabsorption of emitted chlorophyll fluorescence, influence the fluorescence detected from each leaf surface (Cordon and Lagorio, 2007; Terashima and Saeki, 1983; Vilfan et al., 2016; Vogelmann and Evans, 2002). Therefore, differences in absolute fluorescence values and derived fluorescence parameters between the two leaf surfaces can be expected.

In the present study, Y(II) values measured from the adaxial and abaxial surfaces showed some differences in absolute values. Nevertheless, a strong positive correlation was observed between measurements from the two surfaces, and abaxial measurements similarly detected changes in Y(II) during chilling and high-light treatments. A strong correlation between Y(II) from the two surfaces was also observed in strawberry leaves. These results indicate that abaxial measurements can reliably track relative changes in photosynthetic performance, although caution is required when comparing absolute values between the two leaf surfaces. Because leaf anatomy and optical properties vary considerably among plant species, the relationship between adaxial and abaxial fluorescence may also be species dependent and should be considered when applying the Sensor-PAM to different plant species.

Importantly, abaxial measurements provide a practical advantage for a wearable sensor. Attaching the sensor to the abaxial surface allows chlorophyll fluorescence to be monitored continuously at a fixed position without interfering with incident light on the adaxial surface. Thus, rather than providing absolute values directly interchangeable with conventional adaxial PAM measurements, the Sensor-PAM is particularly suited for tracking temporal changes in photosynthetic performance while preserving the natural light environment of the leaf.

### 4.3 Measurement performance of the Sensor-PAM

The Sensor-PAM showed strong correlations with the Reference PAM fluorometer across Arabidopsis, strawberry, and 101 leaves collected from diverse plant species under field conditions. These results indicate that the Sensor-PAM can reliably detect relative differences and changes in Y(II) across different species and environmental conditions. However, regression slopes of less than 1 were observed in the comparisons with Arabidopsis and strawberry under low-temperature and high-light treatments (Fig. 4B, D), suggesting a difference in the quantitative response of Y(II) between the two instruments.

Bland–Altman analysis revealed a systematic underestimation of Y(II) by approximately 0.12 relative to the commercial instrument. The systematic offset and the difference in quantitative response may arise from differences in the optical characteristics of the two instruments, including excitation intensity, optical geometry, and the spectral characteristics of fluorescence detection (Kalaji et al., 2017; Murchie and Lawson, 2013). In particular, the Reference PAM used in this study detects fluorescence at wavelengths above 630 nm using a long-pass filter, whereas the Sensor-PAM selectively detects fluorescence at 680 nm. This difference in detection wavelength may contribute to differences in the measured fluorescence signals between the two instruments.

Importantly, the objective of the Sensor-PAM is not to reproduce exactly the same absolute Y(II) values as a commercial PAM fluorometer, but to reliably monitor temporal changes in photosynthetic performance. The consistently strong correlations observed under both controlled and natural conditions indicate that the Sensor-PAM is suitable for tracking relative changes in Y(II) during long-term continuous measurements. Nevertheless, the systematic offset and differences in quantitative response between the instruments should be considered when directly comparing absolute Y(II) values obtained using the Sensor-PAM with those obtained using conventional PAM fluorometers.

### 4.4 Future perspectives and applications

The wearable Sensor-PAM developed in this study provides a new platform for continuous monitoring of photosynthesis while attached directly to intact plants. By enabling repeated measurements from the same position on the same leaf, the system opens new opportunities for investigating long-term photosynthetic dynamics that are difficult to capture using conventional PAM fluorometers.

Such measurements have considerable potential for studies of plant responses to environmental fluctuations, field phenotyping, and digital agriculture (Fu et al., 2022; Li et al., 2023). Simultaneous deployment on multiple plants could further facilitate comparisons among genotypes, cultivation practices, and environmental conditions.

At present, the Sensor-PAM measures Fv/Fm and Y(II). Expanding the system to additional chlorophyll fluorescence parameters, such as non-photochemical quenching (NPQ), electron transport rate (ETR), and photochemical quenching (qP), represents an important direction for future development. Further improvements in power consumption, miniaturization, and wireless communication will also enable fully autonomous long-term monitoring (Kohzuma and Miyamoto 2024), extending the capabilities and applications of the Sensor-PAM.

## 5 Conclusion

In this study, we developed Sensor-PAM, a wearable chlorophyll fluorescence measurement system capable of continuously monitoring photosynthetic dynamics while attached directly to intact leaves. The system combines a commercially available color sensor with a simple optical design to achieve compact and low-cost PAM measurements. Measurements from the abaxial leaf surface reliably tracked changes in photosynthetic performance and showed strong correlations with those obtained using a commercial PAM fluorometer. Moreover, the Sensor-PAM enabled long-term continuous monitoring under natural environmental conditions. Together, these findings demonstrate the potential of wearable sensing to transform PAM fluorometry from point-based measurements into continuous monitoring of photosynthetic dynamics, providing new opportunities for plant physiology, field phenotyping, and digital agriculture.

## Acknowledgements

The authors thank Masago Housing for providing access to the site used for field measurements.

## CRediT authorship contribution statement

Ko-ichiro Miyamoto: Writing – review & editing, Writing – original draft, Visualization, Validation, Supervision, Software, Resources, Project administration, Methodology, Investigation, Funding acquisition, Formal analysis, Data curation, Conceptualization. Kentaro Ifuku: Resources, Supervision, Writing – review & editing. Tatsuo Yoshinobu: Resources, Supervision, Writing – review & editing. Kaori Kohzuma: Writing – review & editing, Writing – original draft, Visualization, Validation, Supervision, Software, Resources, Project administration, Methodology, Investigation, Funding acquisition, Formal analysis, Data curation, Conceptualization.

## Data availability

The data that has been used is confidential.

## Declaration of generative AI and AI-assisted technologies in the manuscript preparation process

During the preparation of this work, the author used ChatGPT (OpenAI) in order to improve the English language and grammar of the manuscript. After using this tool/service, the authors reviewed and edited the content as needed and take full responsibility for the content of the published article.

