## Supplemental Fig. 1S, Fig. 2S and Table S1 for "Development of a Wearable Sensor-PAM for Continuous Monitoring of Photosynthetic Dynamics"

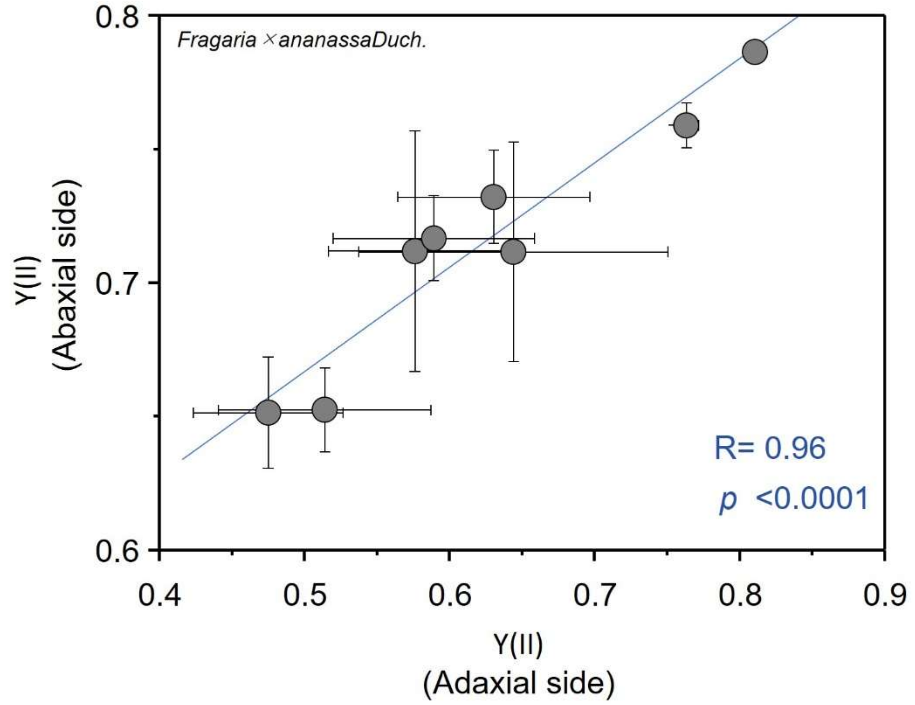


**Fig. S1.** Validation of abaxial chlorophyll fluorescence measurements in strawberry leaves. Correlation between Y(II) values measured from the adaxial and abaxial sides of strawberry (*Fragaria* × *ananassa*) leaves using a commercial PAM fluorometer (Junior-PAM, Walz). Measurements were performed on the same leaves under low-temperature/high-light treatment conditions. The x-axis represents Y(II) measured from the adaxial side, and the y-axis represents Y(II) measured from the abaxial side. Each point represents one treatment condition (n = 8 leaves).

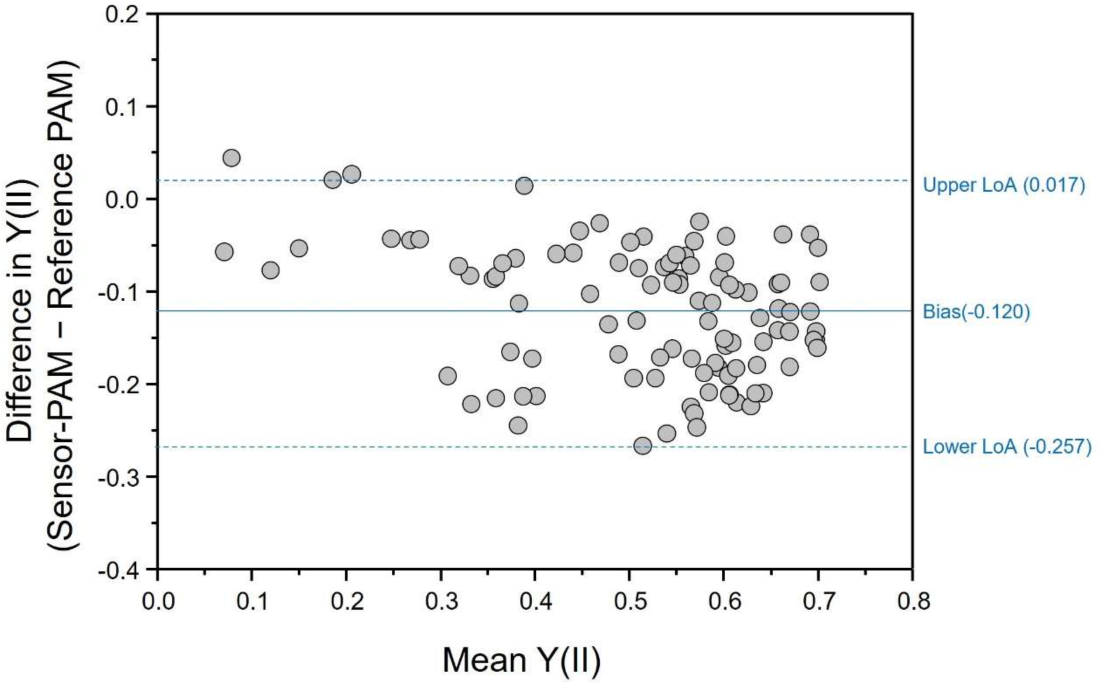


**Fig. S2.** Bland–Altman analysis comparing the Sensor-PAM with the reference PAM fluorometer. Agreement between Y(II) values measured by the Sensor-PAM and the reference PAM fluorometer. The x-axis represents the mean Y(II) of the two instruments, and the y-axis represents the difference between the Sensor-PAM and the reference PAM fluorometer (Sensor-PAM − Reference PAM fluorometer). The solid horizontal line indicates the mean bias (−0.120), and the dashed lines indicate the 95% limits of agreement (−0.257 to 0.017).

**Table. S1. Parts list for the wearable Sensor-PAM. Prices are in USD, as of the time of original manuscript preparation (July, 2026).**

| Section | Part name | Unit Price ($) | Quantity | Total Price ($) |
|---|---|---|---|---|
| Interface | Controller | 11.53 | 1 | 11.53 |
| | FPC connector (SMD type) | 1.33 | 1 | 1.33 |
| | FET (SMD type) | 0.20 | 6 | 1.22 |
| | Voltage regulator (SMD type) | 0.84 | 1 | 0.84 |
| | Custom PCB | 0.48 | 1 | 0.48 |
| | Resister (SMD type) | 0.03 | 14 | 0.43 |
| | Electrolytic capacitor | 0.24 | 1 | 0.24 |
| | Pin header | 0.22 | 1 | 0.22 |
| | Capacitor (SMD type) | 0.10 | 2 | 0.20 |
| Sensor head | Color sensor | 8.23 | 1 | 8.23 |
| | Custom FPC | 2.89 | 1 | 2.89 |
| | Blue LED (SMD type) | 1.22 | 2 | 2.43 |
| | | **Grand total** | **32** | **30.05** |
